# The Electron Transport Chain of *Shewanella oneidensis* MR-1 can Operate Bidirectionally to Enable Microbial Electrosynthesis

**DOI:** 10.1101/2023.08.11.553014

**Authors:** Kathryne C. Ford, Michaela A. TerAvest

## Abstract

Extracellular electron transfer (EET) is a process by which bacterial cells can exchange electrons with a redox active material located outside of the cell. In *Shewanella oneidensis*, this process is natively used to facilitate respiration using extracellular electron acceptors such as Fe(III) or an anode. Previously, it was demonstrated that this process can be used to drive microbial electrosynthesis of 2,3-butanediol (2,3-BDO) in *S. oneidensis* exogenously expressing butanediol dehydrogenase (Bdh). Electrons taken into the cell from a cathode are used to generate NADH, which in turn is used to reduce acetoin to 2,3-BDO via Bdh. However, generating NADH via electron uptake from a cathode is energetically unfavorable, so NADH dehydrogenases couple the reaction to proton motive force. We therefore need to maintain the proton gradient across the membrane to sustain NADH production. This work explores accomplishing this task by bidirectional electron transfer, where electrons provided by the cathode go to both NADH formation and O_2_ reduction by oxidases. We show that oxidases use trace dissolved oxygen in a microaerobic bioelectrical chemical systems (BES), and the translocation of protons across the membrane during O_2_ reduction supports 2,3-BDO generation. Interestingly, this process is inhibited by high levels of dissolved oxygen in this system. In an aerated BES, O_2_ molecules react with the strong reductant (cathode) to form reactive oxygen species, resulting in cell death.

**Importance:** Microbial electrosynthesis is increasingly employed for the generation of specialty chemicals such as biofuels, bioplastics, and cancer therapeutics. For these systems to be viable for industrial scale-up, it is important to understand the energetic requirements of the bacteria to mitigate unnecessary costs. This work demonstrates sustained production of an industrially relevant chemical driven by a cathode. Additionally, it optimizes a previously published system by removing any requirement for phototrophic energy, thereby removing the additional cost of providing a light source. We also demonstrate the severe impact of oxygen intrusion into bioelectrochemical systems, offering insight to future researchers aiming to work in an anaerobic environment. These studies provide insight into both the thermodynamics of electrosynthesis and the importance of bioelectrochemical systems design.

## Introduction

As reliance on fossil fuels becomes increasingly unsustainable from an economic and environmental perspective, researchers work toward alternative energy sources. One solution is microbial electrosynthesis, which is the microbially catalyzed transfer of electrons from an electrode to cells to drive a biochemical reduction reaction(1–3).

Microbial electrosynthesis can be catalyzed by electroactive bacteria capable of interfacing with an electrode surface in a BES or by using the electrode to generate electron carriers, such as H_2_ or formate, that can be taken up by bacteria(4–6). When using electroactive bacteria, a potential is applied to the system to drive oxidation of an electron donor (typically H_2_O) at the anode, and the electrons liberated are taken up by bacteria at the cathode surface. These electrons are used for reduction of a feedstock, such as CO_2_, to the desired product. When CO_2_ is the reactant, microbial electrosynthesis becomes a carbon sink, acting as a carbon-neutral platform to produce biofuels, bioplastics, or specialty chemicals(7, 8). Because CO_2_ is the ideal feedstock for microbial electrosynthesis, much of the existing research focused on bacterial strains or communities with the capacity for autotrophic growth. However, these microbes are inefficient due to slow growth rates and poor interaction with electrodes(9–11). Recent advances in engineering autotrophy raise the possibility of expanding the applications of microbial electrosynthesis beyond the need for native autotrophy or mixed microbial populations(12–15). Therefore, we have focused on a well-understood electroactive bacterial chassis, *Shewanella oneidensis* MR-1, and optimizing product generation^16,17^.

*S. oneidensis* MR-1 is a metal-reducing bacterium with a well-characterized extracellular electron transfer pathway using MtrCAB(16–20). The Mtr pathway allows *S. oneidensis* to respire anaerobically using extracellular, insoluble electron acceptors such as Fe(III) oxides, Mn(IV) oxides, and electrodes(21). As with aerobic respiration, electrons are passed into the quinol pool (menaquinol and ubiquinol) by dehydrogenases in the electron transport chain that oxidizes metabolites such as lactate, formate, and NADH. For extracellular electron transfer, the reduced quinones (quinols) are oxidized by the inner membrane bound cytochrome CymA(22–25). This protein acts as an electron hub, depositing electrons onto periplasmic electron carriers, such as fumarate reductase (FccA) and a small tetraheme cytochrome protein (CctA), to be shuttled onto terminal oxidoreductases. During respiration with an extracellular electron acceptor such as an anode, electrons are transferred to the Mtr pathway, a three-protein complex that spans the outer membrane and extends into the extracellular space(26, 27).

Importantly, the Mtr pathway is reversible, allowing inward electron transfer from a cathode into the cell(2, 26, 28–32). Electrons that are taken up via the Mtr pathway reduce respiratory quinones, and the cell can use the quinols as the electron donor for NAD^+^ reduction by reversing NADH dehydrogenases. NADH can be used to drive a wide variety of intracellular reduction reactions. As a proof-of-concept, we previously demonstrated that reducing power from the electrode can be used to drive the NADH-dependent reduction of acetoin to 2,3-BDO via the heterologous enzyme butanediol dehydrogenase (Bdh) (**Figure 1**). By measuring the accumulation of 2,3-BDO in the system, we can assess the rate and efficiency of electron uptake(28, 33).

**Figure 1.**
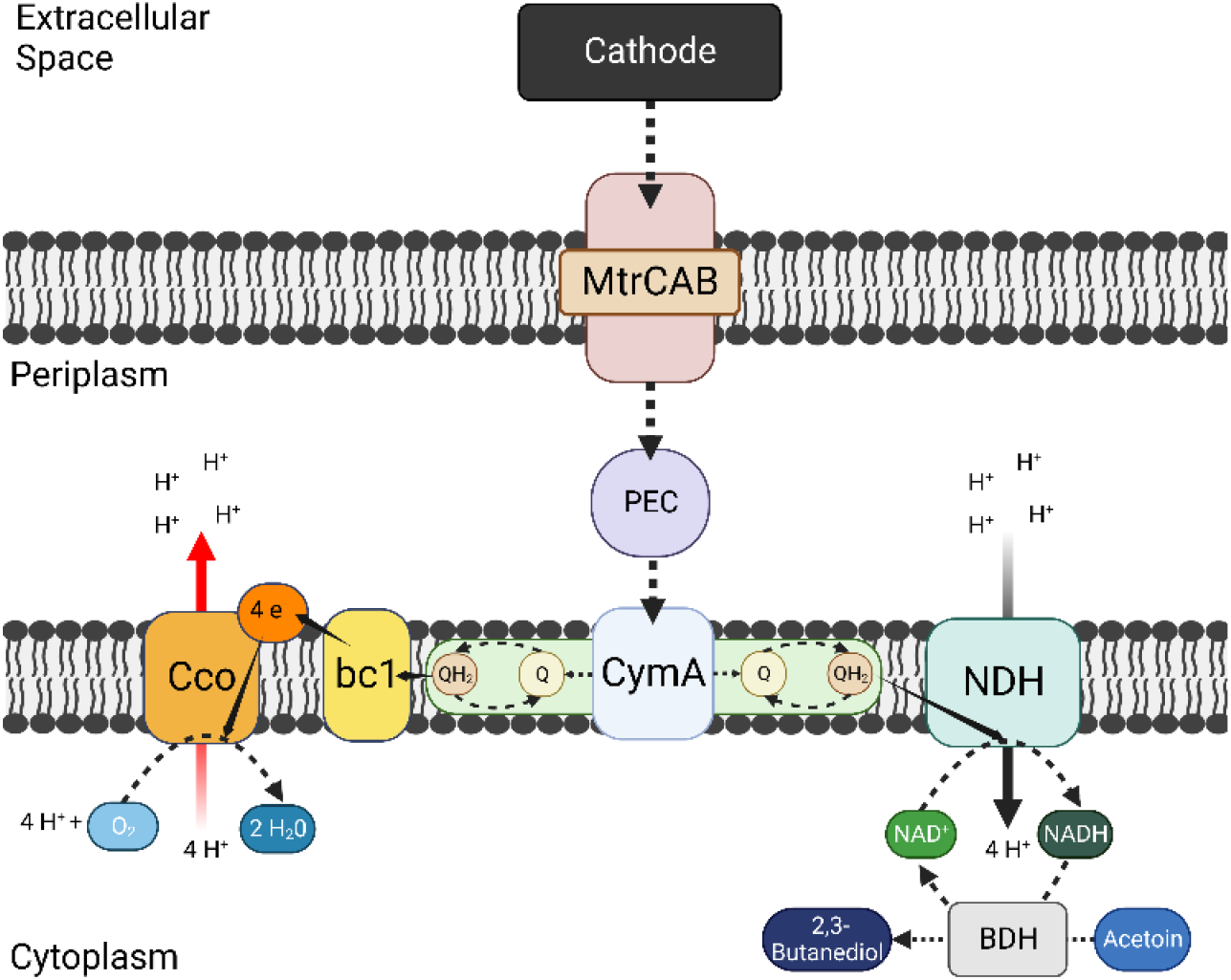
Overview of Bidirectional Electron Transfer to 2,3-BDO. *S. oneidensis* cells are incubated on a cathode poised at -500 mV_Ag/AgCl_. Electrons are taken up by the cell via outer membrane MtrCAB pathway and passed to the inner membrane quinols via various periplasmic electron carriers (FccA, CctA) and CymA. Electrons from the quinol pool are used to reduce NAD^+^ to NADH, catalyzed by NADH dehydrogenases (NDHs) coupling the reaction to proton movement across the inner membrane. The electrode produced NADH is used to reduce exogenous acetoin to 2,3-BDO via butanediol dehydrogenase (BDH).

Inward electron transfer requires a continuous supply of electrons and an electron sink. In this system, acetoin is provided as the electron sink and electrons are supplied by a cathode poised at -0.5 V_Ag/AgCl_. At this electrode potential, electron transfer from the cathode to the MtrCAB complex is thermodynamically favorable. All subsequent reactions in the pathway from electrode to menaquinol are also freely reversible. However, there is a significant energy barrier for electron transfer from menaquinol (-80 mV_SHE_) to form NADH (-320 mV_SHE_) due to the large difference in reduction potential. To overcome the energy barrier, the reaction is catalyzed by ion-coupled NADH dehydrogenases working in reverse. *S. oneidensis* uses both H^+^-coupled and Na^+^-coupled NADH dehydrogenases. In the forward direction, these enzymes couple NAD^+^ reduction to the movement of ions down the electrochemical (Δψ) and proton or sodium gradient (ΔpH or Δ[Na^+^]) known as proton or sodium motive force (PMF or SMF) across the inner membrane(34). The movement of ions down these gradients into the cytoplasm provides the energy needed to power unfavorable chemical reactions; examples of this include formation of ATP via F_O_F_1_-ATP synthase or, as in this case, NAD^+^ reduction by NADH dehydrogenases. In nature, reverse NADH dehydrogenase activity is a means to prevent the potentially lethal overreduction of the quinol pool by generating NADH(25, 32, 34–36).

To enable electron uptake from the electrode, *S. oneidensis* cells are incubated in a BES in the absence of an organic substrate or native terminal electron acceptor. Under these conditions, the cells do not generate PMF via NADH dehydrogenases (Nuo or Nqr) or succinate dehydrogenase (Sdh) because no substrate for these complexes is available(34, 35). Similarly, the absence of a terminal electron acceptor prevents forward electron transport chain flux. Because the reduction of NAD^+^ to NADH requires the free energy provided by PMF, continuous PMF regeneration is necessary for the sustained production of 2,3-BDO. This concept is supported by the prior observation that addition of CCCP (carbonyl cyanide m-chlorophenyl hydrazone), an ionophore that dissipates PMF, results in the cessation of 2,3-BDO production(33). This result underscores an important consideration for microbial electrosynthesis design; how the cell will maintain PMF to continuously drive the reduction reaction forward. To maintain PMF in previous experiments, we introduced proteorhodopsin (PR). PR is a light-driven proton pump that moves protons against the proton gradient into the periplasm, sustaining PMF. Active PR and illumination resulted in an increase of both 2,3-BDO production and cathodic current(28). However, relying on PR as a source of PMF is not a viable solution for scale-up due to well-known issues with light penetration in industrial photobioreactors, and the additional energy cost associated with continuous illumination(28). Understanding this, we sought to utilize bidirectional electron transfer so electrons from the electrode are used for generating both NADH and PMF.

In bidirectional electron transfer, electrons taken up by the cell go towards both the generation of NADH and the reduction of a terminal electron acceptor, e.g., oxygen. This coupling of electron uptake to oxygen reduction by terminal oxidases for PMF generation was first described in *S. oneidensis* by Rowe et al.(32). They found that under carbon-starvation and aerobic conditions, *S. oneidensis* on a cathode generated “non-growth-linked energy” in the form of PMF via terminal oxidase activity. Evidence indicated that PMF was used for production of ATP and reduced cytoplasmic electron carriers (FMNH_2_, NAD(P)H). However, it is still unknown whether this process could continuously generate reducing power for use in MES as there was no sink for NADH. By implementing the Bdh-based system, we sought to determine if bidirectional electron transfer could sustain PMF generation to drive 2,3-BDO generation.

## Results and Discussion

### Eliminating electrode-independent 2,3-BDO Production

We previously demonstrated that the combination of an electrode and active PR led to higher levels of 2,3-BDO production than without either of these components. Before exploring the possibility of using bidirectional electron transfer instead of PR to drive PMF generation, we reexamined 2,3-BDO production in wild-type (WT) *S. oneidensis* MR-1 with and without active PR. Importantly, this experiment was done using an updated version of the previously described experiment protocol. In previous experiments, 35-50% of 2,3-BDO production was independent of the electrode, likely generated using NADH from organic carbon oxidation. To address the high background, we altered the protocol to promote residual organic carbon in the presence of an electron acceptor (anode) to reduce the availability of alternative sources of NADH. Briefly, in the updated protocol *S. oneidensis* is pre-grown in minimal medium (M5) with 20 mM lactate aerobically (or anaerobically with 40 mM fumarate as described later) for 18 hours, followed by inoculation into the BES under aerobic and anodic conditions (+0.2 V_Ag/AgCl_). After six hours, N_2_ sparging is started to switch the cells from using oxygen to the anode as the electron acceptor. In the current study, this anaerobic, anodic phase continued for 40 hours (versus 18 hours in the previous protocol) before the potential is switched to cathodic (-0.5 V_Ag/AgCl_). After this modification, we observed elimination of electrode-independent butanediol production (**Figure 2**, No Potential).

**Figure 2.**
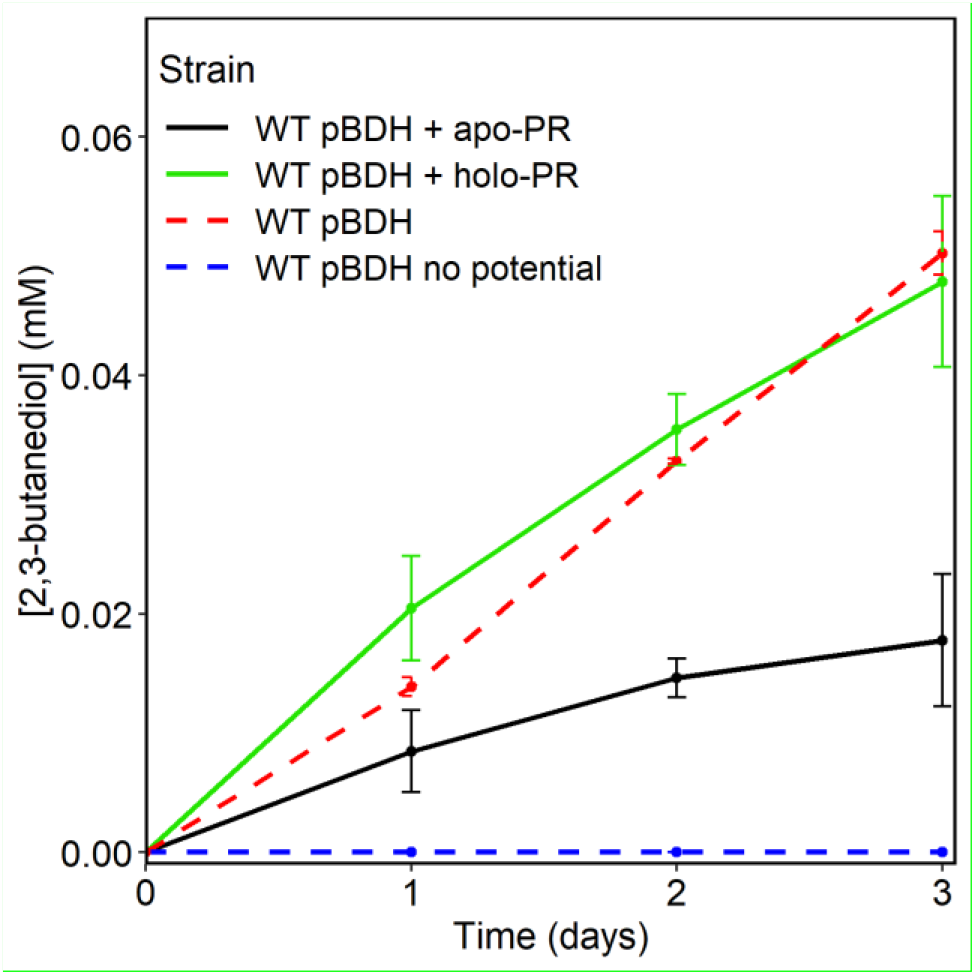
All 2,3-Butanediol Production is Electrode Dependent. Measurement of 2,3-butanediol in BES experiment with modified protocol. WT cells with pBDH or pBDH-PR with (holo-) or without (apo-) retinal as a cofactor, were pre-grown aerobically, washed, and inoculated into anodic BES. After 40 hours potential was switched to cathodic, and acetoin was added to a final concentration of 1 mM (T=0). Samples were collected for HPLC analysis every 24 hours. Lines and error bars represent averages and standard error (n=3).

We tested the amended protocol using a strain expressing PR, with and without the essential cofactor all-*trans*-retinal. Cells with active PR (holo-PR) produced more 2,3-BDO than those with inactive PR (apo-PR). Interestingly, cells with active PR produced approximately the same amount of 2,3-BDO as cells not expressing PR, while cells with inactive PR showed a decrease in 2,3-BDO production (**Figure 2**). This finding suggests that while PMF generation by PR supports an increase in 2,3-BDO production, the metabolic burden or membrane occupancy constraints of expressing PR outweigh the benefits. Moreover, this result suggests that there is an unaccounted-for source of PMF in the absence of PR. Another source of PMF appears more likely than the possibility that PMF is unnecessary, based on experimental evidence and thermodynamic calculations. Our prior work demonstrated that 2,3-BDO production is halted when PMF is dissipated by CCCP(33). Additionally, electron transfer from quinols to form NADH cannot occur at an appreciable rate without the energy gained from proton translocation across the membrane. To sustain the NADH dehydrogenase-catalyzed reaction, which utilizes PMF, there must be a mechanism to replenish the proton gradient. We considered formate dehydrogenase and F_o_F_1_ ATP synthase as possible PMF sources, but found them unlikely due to the lack of a formate or ATP source. Therefore, we speculated that trace amounts of oxygen entering the BES could be sufficient to enable bidirectional electron transfer.

### Bidirectional Electron Transfer to Oxygen and NAD^+^

We investigated the possibility of bidirectional electron transfer as the unknown source of PMF because this reaction could be powered by the electrode, and the substrate (O_2_) is readily accessible. Although the working chamber was continuously degassed by N_2_ bubbling (99.999% N_2_), we suspected that the environment was microaerobic. The BESs may not be completely airtight due to the use of neoprene tubing and plastic connectors, and the possibility of oxygen diffusion from the anode through the ion exchange membrane (**Supp. Figure S1**). Additionally, the N_2_ tank used can contain up to 1 ppm O_2_ contamination, per the product specifications (Airgas). To ascertain if oxygen was present, we inserted an optical dissolved oxygen (DO) probe into the BES and conducted an experiment as normal. The DO in the working chamber was at ∼100% saturation before inoculation, decreased to ∼60% saturation upon cell addition, and dropped to ∼1% upon N_2_ bubbling (**Figure 3A**). This single experiment produced 0.046 mM 2,3-BDO over 3 days, which is consistent with previous experiments (Figure 2.3C).

**Figure 3.**
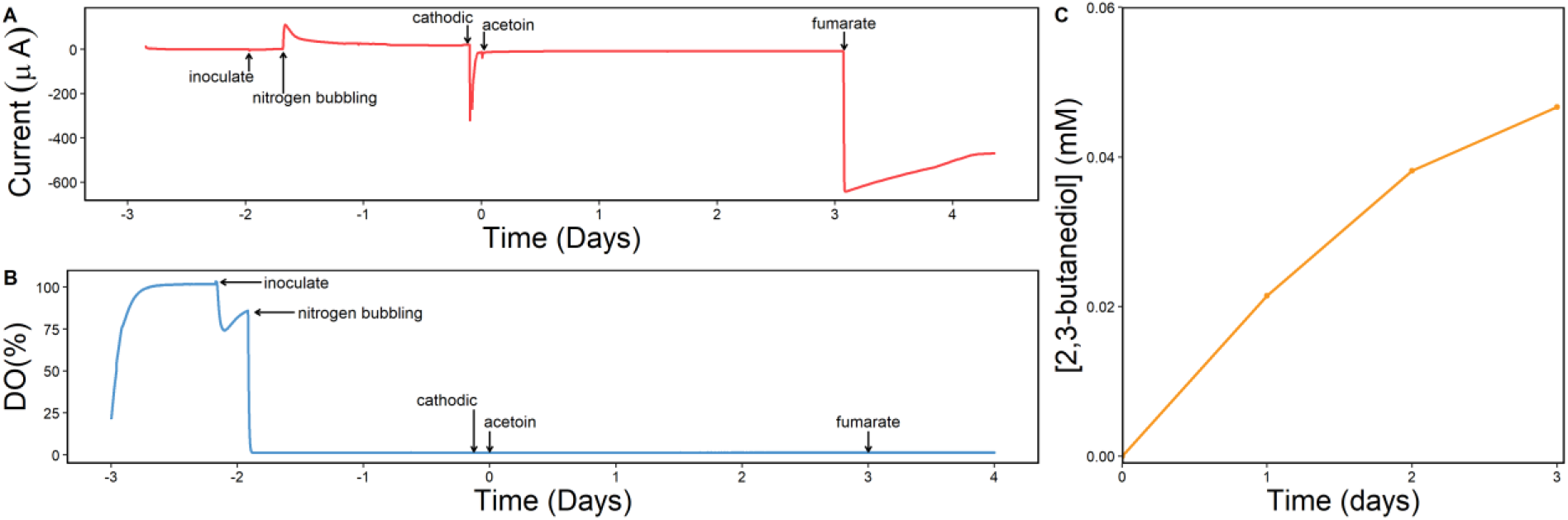
Presence of Oxygen in BES. Time course of the experiment from setup (Day -3) to final timepoint (Day 4), showing the current (A) and dissolved oxygen (B). Samples were taken daily from Day 0 to Day 3 to quantify 2,3-BDO production (C) (n=1).

Production of one 2,3-BDO molecule from acetoin requires oxidation of one NADH, which in turn depletes four H^+^ from available PMF via Nuo. One molecule of oxygen (O_2_) allows translocation of four H^+^ across the membrane if it is reduced by either of the proton-pumping terminal oxidases, Cco and Cox(37–39). Therefore, the reduction of one O_2_ molecule can sustain production of one molecule of 2,3-BDO from the perspective of PMF balance. The 1% saturation DO concentration observed is equivalent to 0.073 mg/L (30°C, 856 elevation), or ∼2 µM of oxygen available throughout the experiment(40). Considering this concentration, there is more than enough oxygen available to support 0.046 mM 2,3-BDO production over 72 hours (0.638 µM 2,3-BDO/hour) as the sole source of PMF.

To verify the contribution of terminal oxidases in generating PMF during electron uptake, we compared current and 2,3-BDO production between WT MR-1 and a strain lacking all 3 terminal oxidases (Δ*cyd*Δ*cco*Δ*cox*, here named Δ*oxidase*)(32). This strain cannot use O_2_ as a terminal electron acceptor meaning that even with trace amounts of oxygen, the mutant cells would not be able to use it.(41) This mutant cannot grow in aerobic conditions, so further comparisons were performed using anaerobic pre-growth of all strains. To ensure consistency, we compared anaerobic growth of WT versus Δ*oxidase* cells in M5 minimal medium (20 mM lactate, 40 mM fumarate). We observed similar growth rates (**Supp. Figure S2**), so for MES experiments, these strains were pre-grown anaerobically. WT cells pre-grown in an anoxic environment produced 0.047 ± 0.002 mM butanediol, consistent with previous work, and exhibited similar current profiles and magnitudes (**Figure 4**). Conversely, the Δ*oxidase* strain produced minimal butanediol, with only a small amount accumulating by day 6, and less than half the current of WT cells. This result highlights that the cells were unable to sustain electrode-dependent acetoin reduction in the absence of aerobic terminal oxidases.

**Figure 4.**
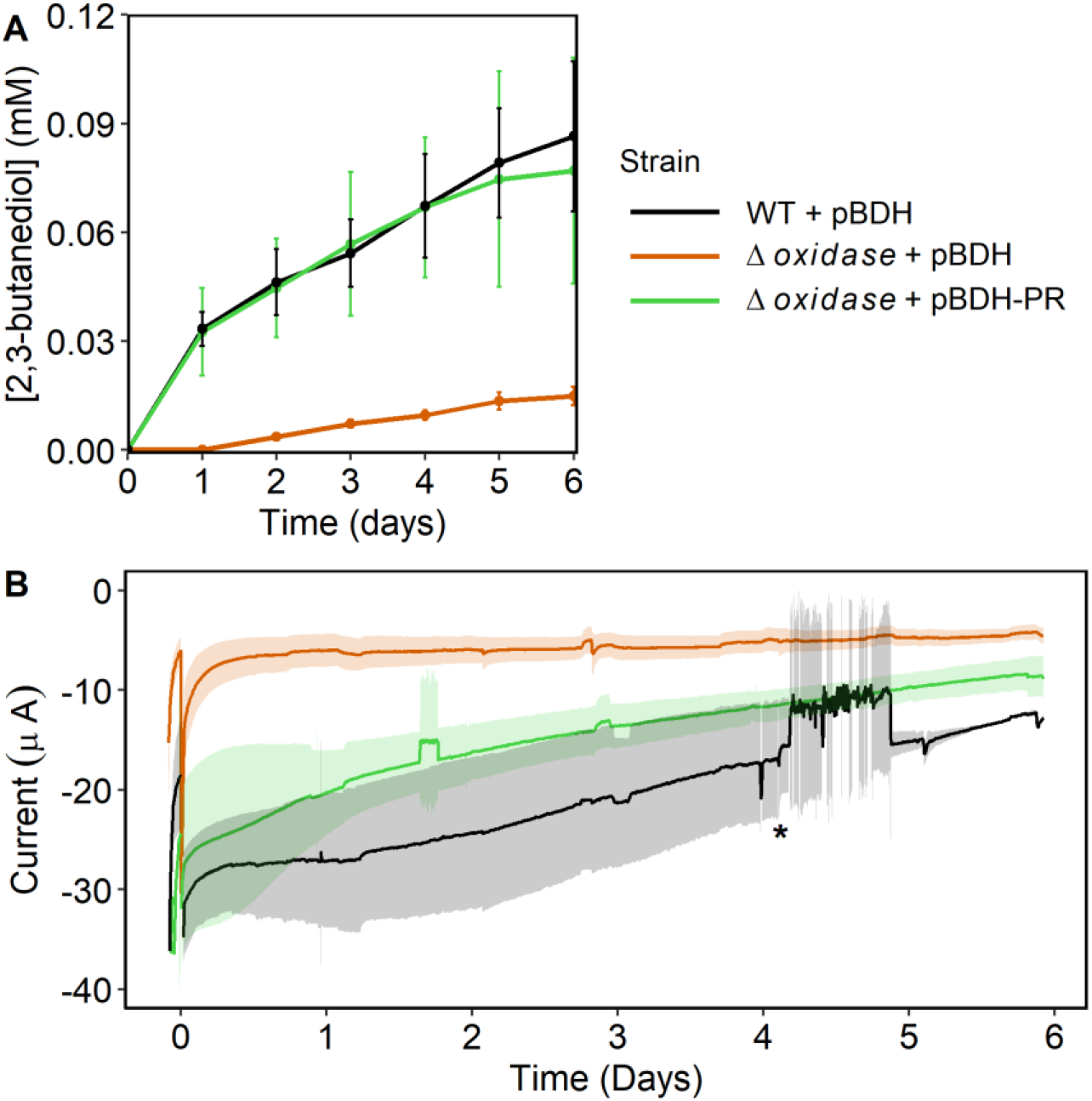
2,3-BDO and Current Production in Microaerobic BES. (A) 2,3-Butanediol accumulation in BES with WT pBDH, Δ*oxidase* pBDH, and Δ*oxidase* pBDH-PR with nitrogen bubbling. (B) Cathodic current from BES. Noisy current marked by (*) was due to stirring issues with one of the BES replicates. Points (A) and lines (B) represent averages with standard error bars, n=3.

To confirm that the observed phenotype was due to the loss of PMF from proton-pumping oxidase activity, the Δ*oxidase* strain was functionally complemented by PR expression. If 2,3-BDO production is rescued by PR, it indicates that the loss of 2,3-BDO generation in Δ*oxidase* is caused by a loss of PMF as opposed to off target effects, such as changes in gene regulation. When PR was expressed in Δ*oxidase*, we observed a restoration of 2,3-BDO production and partial rescue of current (**Figure 4**). In this instance, electron transfer to form NADH but not O_2_ is restored, and as one O_2_ is required to produce one 2,3-BDO molecule, a 50% rescue of current is consistent with our model. Taken together, these results support the hypothesis that PMF is a limiting resource, and the proton pumping activity of oxidases in this microaerobic environment is essential to continuous electron transfer to form NADH.

### Reactive Oxygen Species Formation

We next investigated whether increasing DO in the BES, and by extension oxidase activity, would result in an increase in 2,3-BDO. To do this, BESs were not sparged with N_2_ to allow passive aeration. DO measurements indicated a highly oxygenated environment (300 μM) in the BES (**Supp. Figure S3**). This condition resulted in a severe decrease in 2,3-BDO production relative to the N_2_-bubbling microaerobic condition (**Figure 5**). Current was greatly inflated by O_2_ intrusion (data not shown).

**Figure 5.**
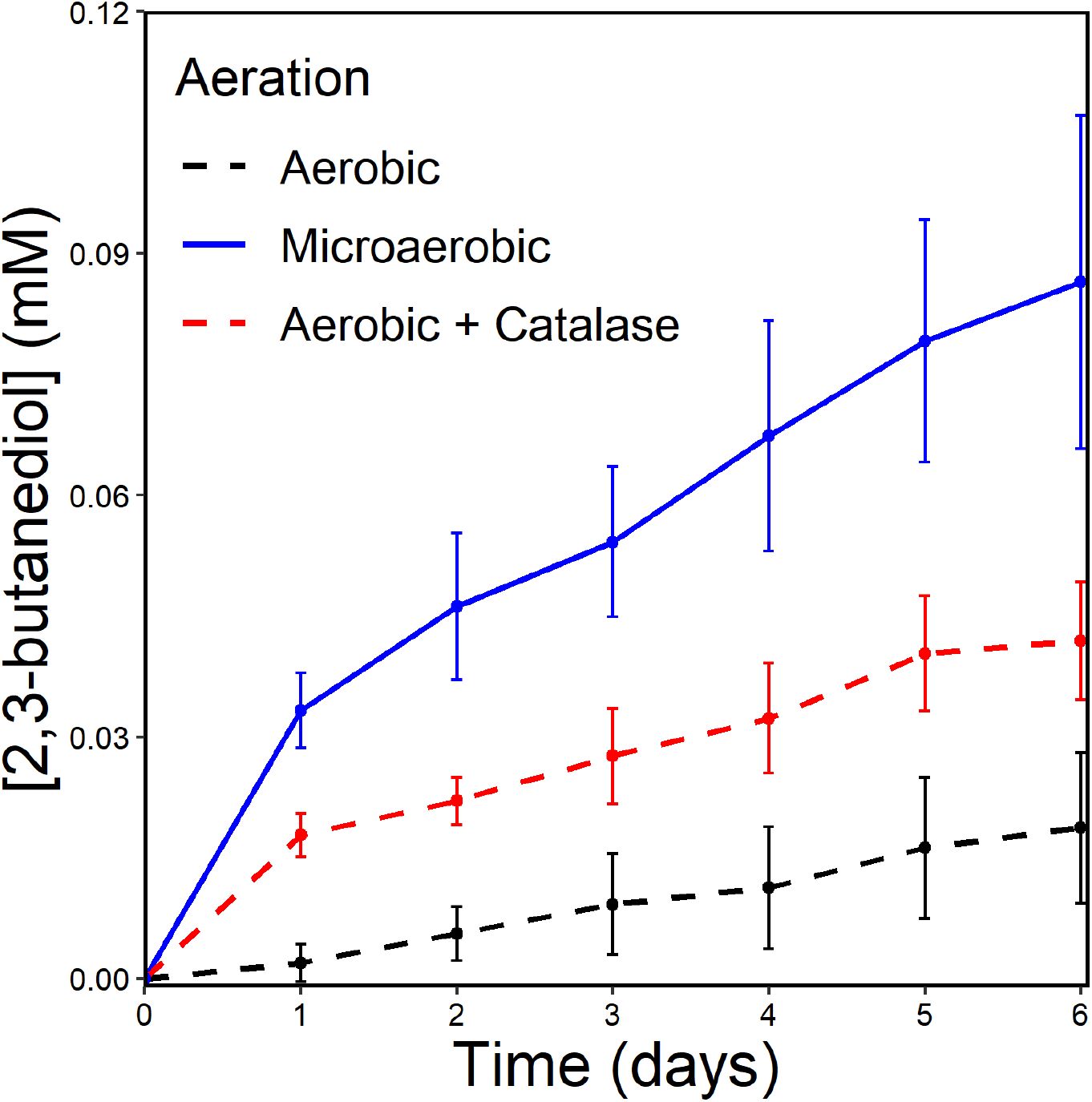
2,3-BDO Production in Aerobic and Microaerobic BES. (A) 2,3-Butanediol accumulation in BES with WT pBDH with N_2_ bubbling (Microaerobic), and passive aeration (Aerobic) with or without the addition of 0.3 U/mL catalase. Points represent averages with standard error bars, n=3.

The failure of increased DO to translate to an increase in 2,3-BDO accumulation could be attributed to three factors: decreased *mtrCAB* expression, formation of reactive oxygen species (ROS), or a shift in electron flow to favor oxygen reduction over NADH generation. In the presence of oxygen, *S. oneidensis* MR-1 decreases expression of anaerobic respiration pathways such as Mtr in favor of aerobic respiration and a decrease in Mtr expression will likely result in a decrease in inward ET(42). However, the cells are not actively growing under the experimental conditions, and it is improbable that a significant shift in the proteome occurred under the carbon starvation conditions of the experiment. Similarly, while loss of all electron flux in favor of oxygen reduction is possible, the small amount of 2,3-BDO produced during passive aeration suggests there are still electrons going towards NADH formation. Additionally, it has been shown that bidirectional electron transfer to NAD^+^ and oxygen occurs under active aeration; if all electrons were being lost to oxygen, the effect would likely have been more pronounced under those conditions(32). Further research done to optimize DO concentration should explore this possibility.

To assess the possibility of ROS formation, we measured hydrogen peroxide (H_2_O_2_) in the BESs. In the presence of oxygen and a strong reductant, such as a cathode or reduced flavin, O_2_ can be reduced to form H_2_O_2_(42–45). H_2_O_2_ accumulation in the BESs may result in cell death and a decrease in the ability to produce 2,3-BDO. H_2_O_2_ can also react directly with 2,3-BDO, possibly leading to reduced accumulation because of abiotic degradation(46, 47). To investigate if ROS accumulated in the passive aerobic condition, experiments with WT pBDH were performed with and without N_2_ bubbling, with and without potential, and samples were taken for colony forming unit (CFU) and H_2_O_2_ measurements. When the potential was swapped from anodic to cathodic, we observed an immediate drop in CFUs/mL and generation of H_2_O_2_ in BES with passive aeration, while the microaerobic BES maintained the same levels of both (Figure 2.6). There was no detectable peroxide formation in the no potential controls (data not shown). This result demonstrates that the formation of H_2_O_2_ is dependent on the presence of oxygen and a cathode. The formation of H_2_O_2_ was correlated with ∼2.5 log_10_ cell death in the first 3 hours.

We next explored whether addition of catalase (an H_2_O_2_ degrading enzyme) could reduce H_2_O_2_ accumulation. This approach has the potential to harness the benefits of oxygen inclusion, such as PMF generation, while minimizing the production of harmful by-products(42). Aerobic BESs were run with the addition of 0.3 U/mL catalase added immediately before the potential was switched to -0.5 V_Ag/AgCl_. Aerobic BESs with catalase did not result in the same rapid decrease in CFUs/mL is increase in H_2_O_2_ as those without, as well as a less prominent spike in peroxide formation (**Figure 6**). Catalase addition also resulted in a partial rescue of 2,3-BDO production (**Figure 5**). These results show that one of the challenges with oxygen inclusion in a BES is the cytotoxic formation of H_2_O_2_, resulting in cell death and decrease in product yield. Future experimentation with aerated BES should focus on optimizing DO concentration to maximize reverse electron flux to oxygen reduction relative to forward electron flux to NAD^+^; having high oxidase activity to generate PMF without losing electron flow to NADH formation.

**Figure 6.**
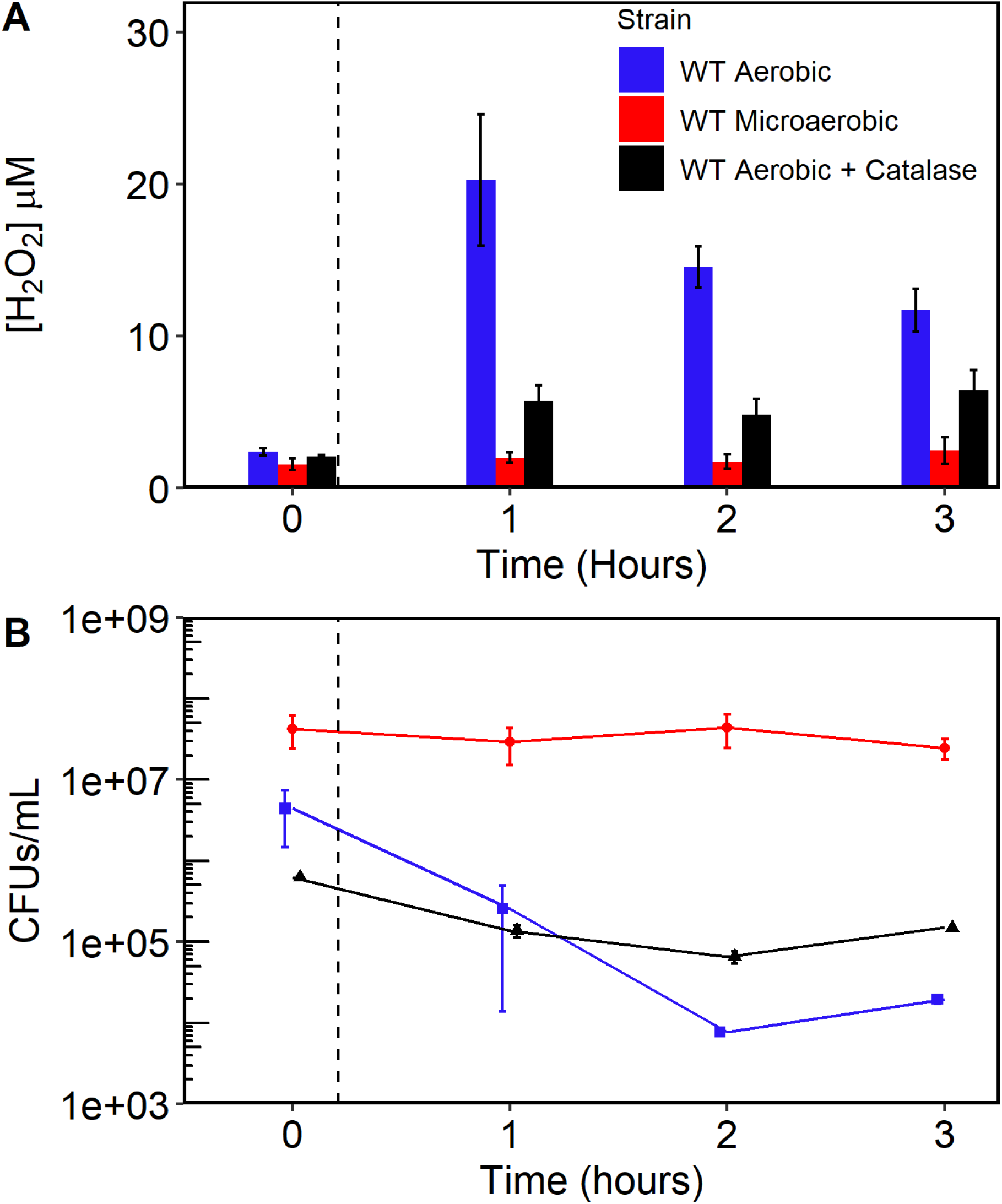
CFUs and [H_2_O_2_] in Aerobic and Microaerobic BES. (A) CFUs/mL and (B) [H_2_O_2_] µM in bulk medium of working chamber. Dashed line represents the potential change from 0.2 V_Ag/AgCl_ to -0.5 V_Ag/AgCl_. Points represent averages of n=3 with standard error bars. Lines are included in CFU/mL data to guide the eye. CFUs/mL at inoculation (∼46 hrs. prior to T=0) for all conditions were ∼1.8 × 10^8^.

## Conclusion

Effective microbial electrosynthesis requires attention to detail in both BES design and bacterial physiology. In the system discussed here, understanding the thermodynamic factors involved in driving inward electron transfer is crucial. The reversible nature of the electron transport pathways, which enables cells to use electrodes as electron acceptors and donors, depends on the reduction potential of each step(22, 48–52). Electron transfer reactions from electrode to quinol pool are freely reversible, but the final transfer from menaquinol to NADH formation has a much larger shift in potential between donor (-80 mV) and acceptor (-330 mV). This barrier is overcome by NADH dehydrogenases catalyzing the reaction and coupling the reduction to PMF utilization. In this work, we show that during electron transfer from an electrode to NADH, PMF can be regenerated by bidirectional electron transfer. Importantly, *S. oneidensis’* native aerobic terminal oxidases (Cco, Cox, Cyd) can sustain PMF via oxygen reduction without fully redirecting the flow of electrons away from NADH. This ability was best demonstrated in microaerobic conditions, where the DO concentration struck the balance between electron flow to oxygen and NAD^+^. Higher levels of oxygen had the off-target effect of generating H_2_O_2_ that resulted in cell death. While the conditions tested here were limited to microaerobic and passively aerobic, future work should focus on fine tuning the DO in BES. This could be done through a combination of oxygen scavengers, gas mixing/modulating inflow, inclusion of other ROS neutralizing enzymes such as superoxide dismutase, or selective deletion of native oxidases as they have varying oxygen affinities and proton pumping efficiencies. The goal should be to balance the redox state of the quinone pool to maximize the flow of electrons ‘uphill’ to NAD^+^ relative to the energetically favorable reduction of oxygen. Taken together, this work shows the strong influence even trace oxygen has on the energetics of inward electron transport.

## Materials and Methods

**Table 1.**

| Strain or Plasmid | Description | Source |
| --- | --- | --- |
| ***S. oneidensis*** |  |  |
| MR-1 | Wild type *S. oneidensis* | Meyers and Nealson, 1988 |
| Δoxidase | Mutant with gene deletion of *cco*, *cyd*, *cox*, (SO2361–SO2364, SO3285–SO3286, SO4606–SO4609) | Rowe et al. 2018 |
| **Plasmids** |  |  |
| pBDH | pBBR1MCS2 bearing butanediol dehydrogenase gene from *Enterobacter cloacae*, kan<sup>R</sup> | Tefft and TerAvest, 2019 |
| pBDH-PR | pBBR1MCS2 bearing butanediol dehydrogenase from *Enterobacter cloacae* and proteorhodopsin (uncultured marine gamma proteobacterium EBAC31A08), kan<sup>R</sup> | Tefft and TerAvest, 2019 |

### Strains and Plasmids

Strains and plasmids used are listed in Table 2.1. *S. oneidensis* MR-1 strains were grown at 30 °C and shaking at 275 rpm for aerobic growth, and no shaking for anaerobic growth (∼5% H_2_, balanced with N_2_). For BES experiments, MR-1 was pre-grown aerobically in 5 mL of lysogeny broth (LB) supplemented with 50 μg/mL kanamycin for strains with pBBR1-BDH, for inoculating minimal medium. For pre-growth, cells were grown in M5 minimal medium containing: 1.29 mM K_2_HPO_4_, 1.65 mM KH_2_PO_4_, 7.87 mM NaCl, 1.70 mM NH_4_SO_4_, 475 μM MgSO_4_·7 H_2_O, 10 mM HEPES, 0.01% (w/v) casamino acids, 1× Wolfe’s vitamin solution, and 1× Wolfe’s mineral solution, then the pH adjusted to 7.2 with 5 M NaOH. After autoclaving, D,L-lactate was added to a final concentration of 20 mM. During anaerobic pre-growth, fumarate was added to a final concentration of 40 mM and 400 mL of medium was used per repeat. During bioelectrochemical experiments, the M5 medium recipe was amended to 100 mM HEPES, 0.2 μM riboflavin, and no D,L-lactate, fumarate, or casamino acids.

### Growth Curves

For anaerobic growth experiments, cells were pre-grown in 5 mL LB supplemented with 40 mM fumarate and 20 mM D,L-lactate. Cells from the overnight culture were washed with M5 medium and resuspended to an OD_600_ of 0.05 in 2 mL M5 medium in a 24-well plate. OD_600_ was measured every 15 minutes for 35 hours in an anaerobic plate reader (BioTek, HTX). This protocol was repeated 3 times for replication.

### Bioelectrochemical System Experiments

BES experiments were conducted in custom made two-chamber bioreactors kept at 30°C as described in previous work (Tefft and TerAvest 2019)(28), and a similar set up to work described in (Tefft et al. 2022)(29). The working chamber was filled with 144 mL amended M5 medium, with 0.2 μM riboflavin being added an hour before inoculation, and the counter chamber contained ∼150 mL of 1x PBS. For experiments run with PR, green LED lights were attached to the reactors. Bioreactors were autoclaved for 45 minutes, then connected to a potentiostat (VMP, BioLogic USA) and current data was collected every 1 s for the course of the experiment. After the initial setup, the working electrode poised at an anodic potential of +0.2 V_Ag/AgCl_ for ∼16 hours. For aerobic pre-growth experiments, cells were grown in two 50-mL cultures of M5 in 250-mL flasks for each bioreactor (6 total for 3 replicates) for 18 hours. For anaerobic pre-growth experiments, cells were grown in 400-mL cultures of M5 in 1-L flasks for each bioreactor (3 total for 3 replicates) for 18 hours. For experiments with PR, 400 μL 20 mM all-*trans*-retinal was added after 17 hours of growth as the essential cofactor for PR. Cultures were transferred to a 50-mL conical tube and centrifuged at 8000 rpm (Thermo Scientific ST8R; Rotor: 75005709) for 5 minutes. Pellets were washed twice in 30 mL M5 (100 mM HEPES, no carbon) and then resuspended in M5 (100 mM HEPES, no carbon), to a final OD_600_ of 3.6 in 10 mL. Then, 9 mL of this normalized resuspension was inoculated into the working chamber of the bioreactor using a sterile 10 mL syringe with an 18 g needle. Six hours after inoculation, N_2_ gas (99.999%, AirGas) was bubbled into reactors through a 0.2 μM filter, and a bubbler attached to a 0.2 μM filter connected to the gas outlet. For 40 hours after N_2_ bubbling, reactors were maintained at an anodic potential of +0.2 V_Ag/AgCl_, before being changed to a cathodic potential of -0.5 V_Ag/AgCl_. After three hours at cathodic potential, 17 mL of a sterile, de-gassed 10 mM acetoin solution was added to a final concentration of 1 mM in the bioreactor (Final volume in working chamber = 170 mL). The bioreactors were sampled (2 mL) immediately after acetoin addition for OD_600_ and HPLC analysis every 24 hours for 144 hours.

### DO Measurements

DO measurements shown in Figure 3 were collected using a Hamilton VisiFerm DO sensor and ArcAir Software. The probe was calibrated before each experiment as described in the manual. The probe was inserted into the BES prior to autoclaving and secured with a rubber gasket. DO measurements were recorded every 5 s during the experiment. To ensure that the inclusion of the DO probe did not interfere with oxygen intrusion into the system, we also utilized a smaller fiber optic DO probe and collected data every 30 s using a NeoFox Fluorimeter and Software (Ocean Insight). The probe consists of a patch made from 5% mixture of polymer (poly(2,2,2-trifluoroethyl methacrylate), Scientific Polymer Products Inc.) and 5 mM porphyrin (Pt(II) meso-tetra(pentafluorophenyl)porphine, Frontier Scientific) dissolved in a 50/50 mixture of 1,4-dioxane and 1,2-dichloroethane (Sigma Aldrich). This patch is deposited onto the end of a fiber optic probe(53). Results from this probe corroborated observations made with the Hamilton Probe (Supp. Figure 2.3A). Data from passively aerobic BES (Supp. Figure 3B) was collected using the smaller fiber optic probe.

### CFU Plating

During BES experiments, samples were taken every ∼24 hours starting at inoculation, with additional time points in the three hours following potential change from anodic to cathodic. These samples were used for CFU plating, H_2_O_2_ measurements, and HPLC analysis. Samples were serially diluted in a 96-well plate and 10 μL of each of 8 dilutions (10^0^-10^-7^) was plated on LB + Kan. Dilutions with between ∼10^1^-10^2^ CFUs were counted and back calculated to determine CFUs/mL in bulk solution. Mean and standard error were calculated for biological replicates (n=3).

### H_2_O_2_ Measurements

At each sampled time point, H_2_O_2_ formation was measured using the Pierce™ Quantitative Peroxide Assay Kit (ThermoFisher, Cat: 23280) according to the kit instructions. In brief, 20 μL of sample was mixed with 200 μL of reagent mixture in a 96-well plate, and absorbance was read at 595 nm. Sample values were compared to a standard curve with background subtraction of cell-only controls in 1xPBS to exclude any interference from cell OD_600_. Mean and standard error were calculated for biological replicates (n=3).

### HPLC analysis

HPLC analysis was performed as previously described (Tefft and TerAvest, 2019) with the amendments described in (Tefft et al., 2022)(28, 29). Sample analysis was performed on a Shimadzu 20A HPLC, using an Aminex HPX-87H (BioRad, Hercules, CA) column with a Microguard Cation H^+^ guard column (BioRad, Hercules, CA) at 65 °C with a 0.5 ml/min flow rate. 2,3-butanediol concentration in samples was calculated by comparing sample value to an external standard curve.

### Data analysis

Analysis of HPLC data, DO %, OD, current data, and growth curve data was done using RStudio using the following packages: ggplot2, dplyr, ggpubr, plyr, data.table, stringr, and growthcurver(54–60).

## Author Contributions

K.C.F. and M.T. conceptualized the project. K.C.F. lead the investigation and data visualization under the supervision of M.T. K.C.F. wrote the original draft of the manuscript, with review and edits by M.T.

## Acknowledgements

The authors would like to thank Nathan Frantz (Michigan State University - Department of Chemistry) and Dr. Denis A. Proshlyakov (Michigan State University - Department of Physiology) for their assistance with DO measurements. This research was supported by the National Science Foundation Graduate Research Fellowship Grant No. (DGE-1848739) to K.C.F., and a National Science Foundation CAREER Award (1750785) and 2018 Beckman Young Investigator Award to M.T.

## 2.9 Supplementary Figures

**Supplementary Figure 2.1.**
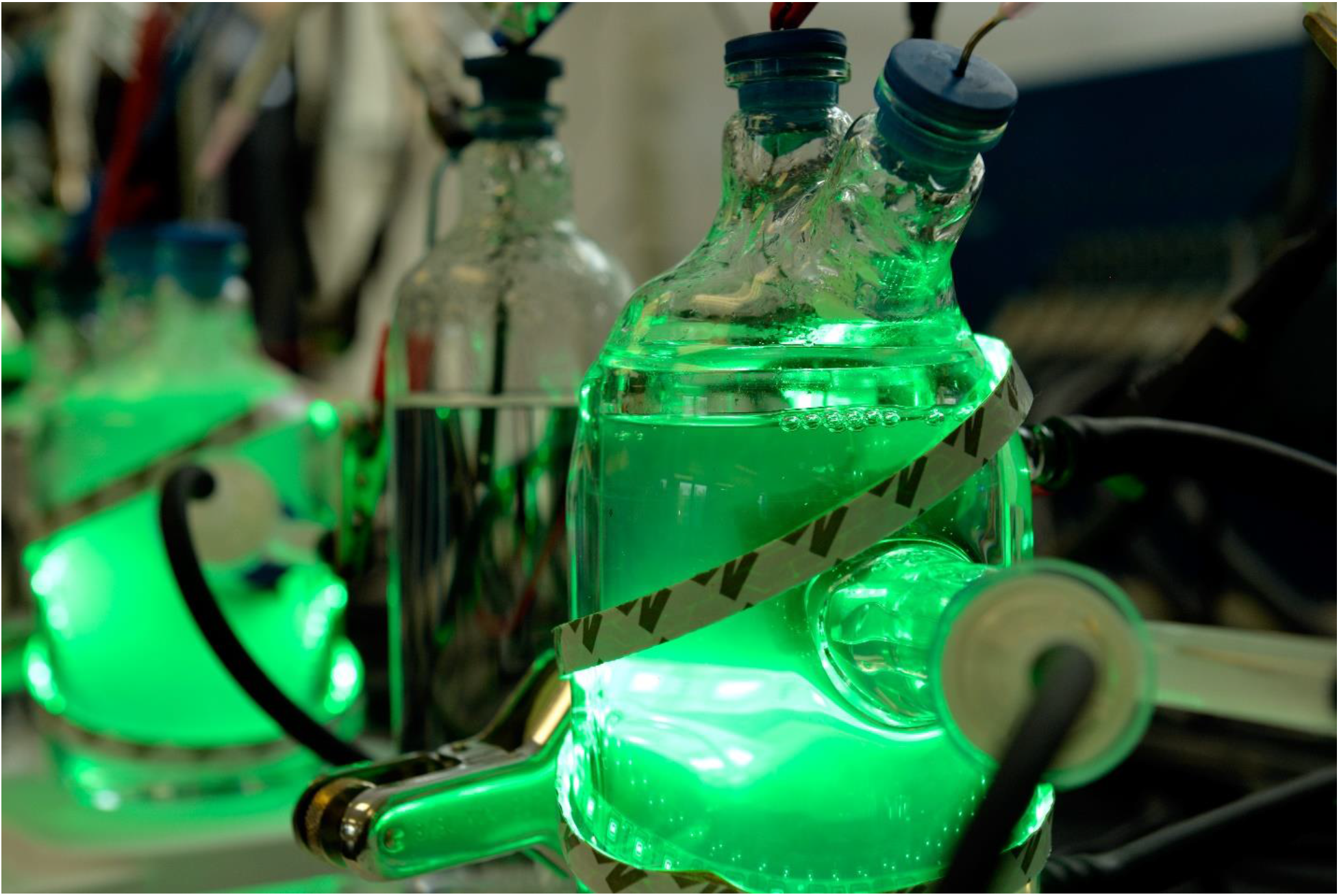
Bioelectrochemical System Design. Setup of our 2-chamber BES during an experiment with cells expressing PR, hence the inclusion of the green lights. Chambers are sealed with blue stoppers and N_2_ is bubbled in from a neoprene tube attached to a 0.2 µm filter (foreground).

**Supplementary Figure 2.2.**
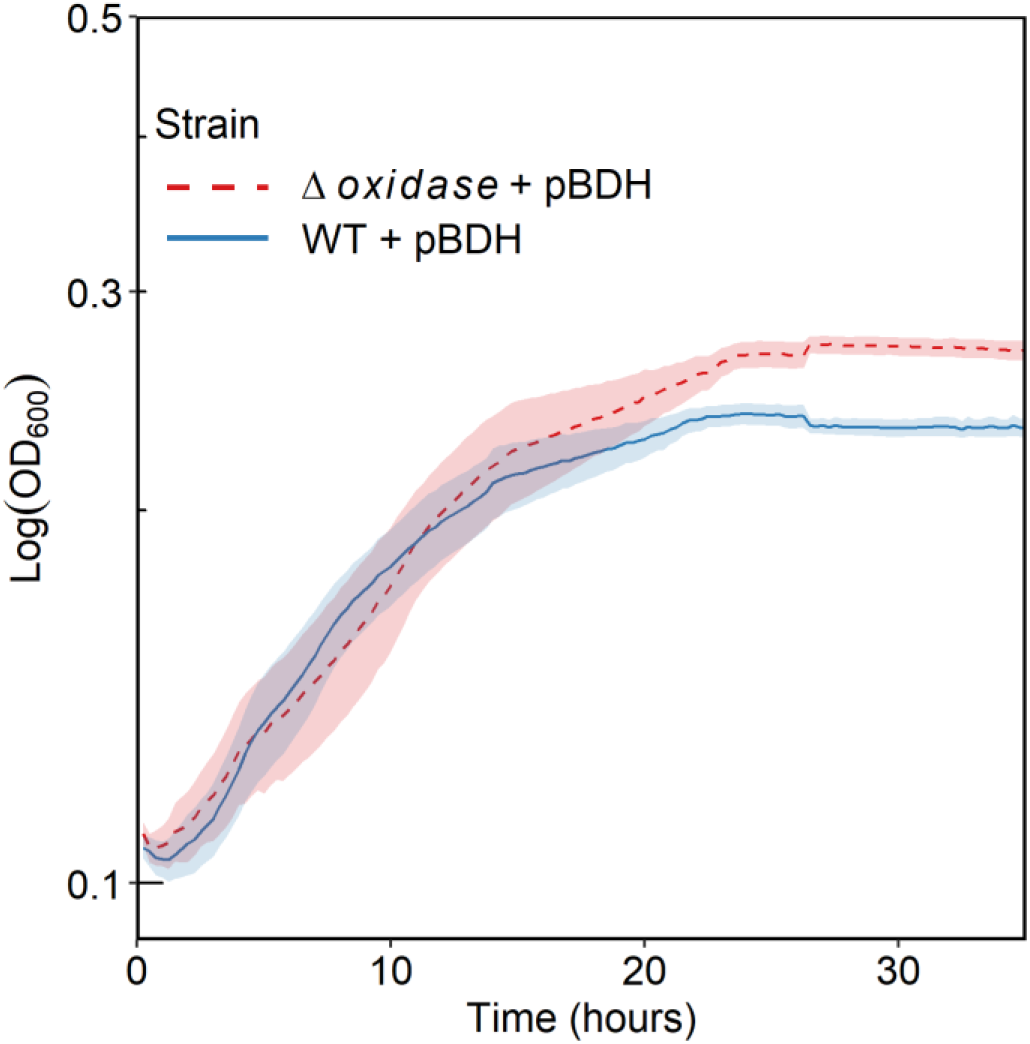
Growth curves of WT and Δ*oxidase*. Each strain was struck on LB + 20 mM _D,L_-lactate + 40 mM fumarate + Kan plates and incubated anaerobically at 30°C. Three single colonies were used to start 5 mL anaerobic overnight cultures in LB + 20 mM _D,L_-lactate + 40 mM fumarate + Kan. Overnight cultures were spun down and pellets resuspended in M5 + 20 mM _D,L_-lactate + 40 mM fumarate + Kan to an OD_600_ of 1.0. These samples were used to inoculate 2 mL of M5 + 20 mM _D,L_-lactate + 40 mM fumarate + Kan in a 24-well plate at an initial OD_600_ of 0.05. OD_600_ was measured every 15 minutes for 35 hours. Lines and shaded region represent the mean and standard error, respectively, for n=3 replication.

**Supplementary Figure 2.3.**
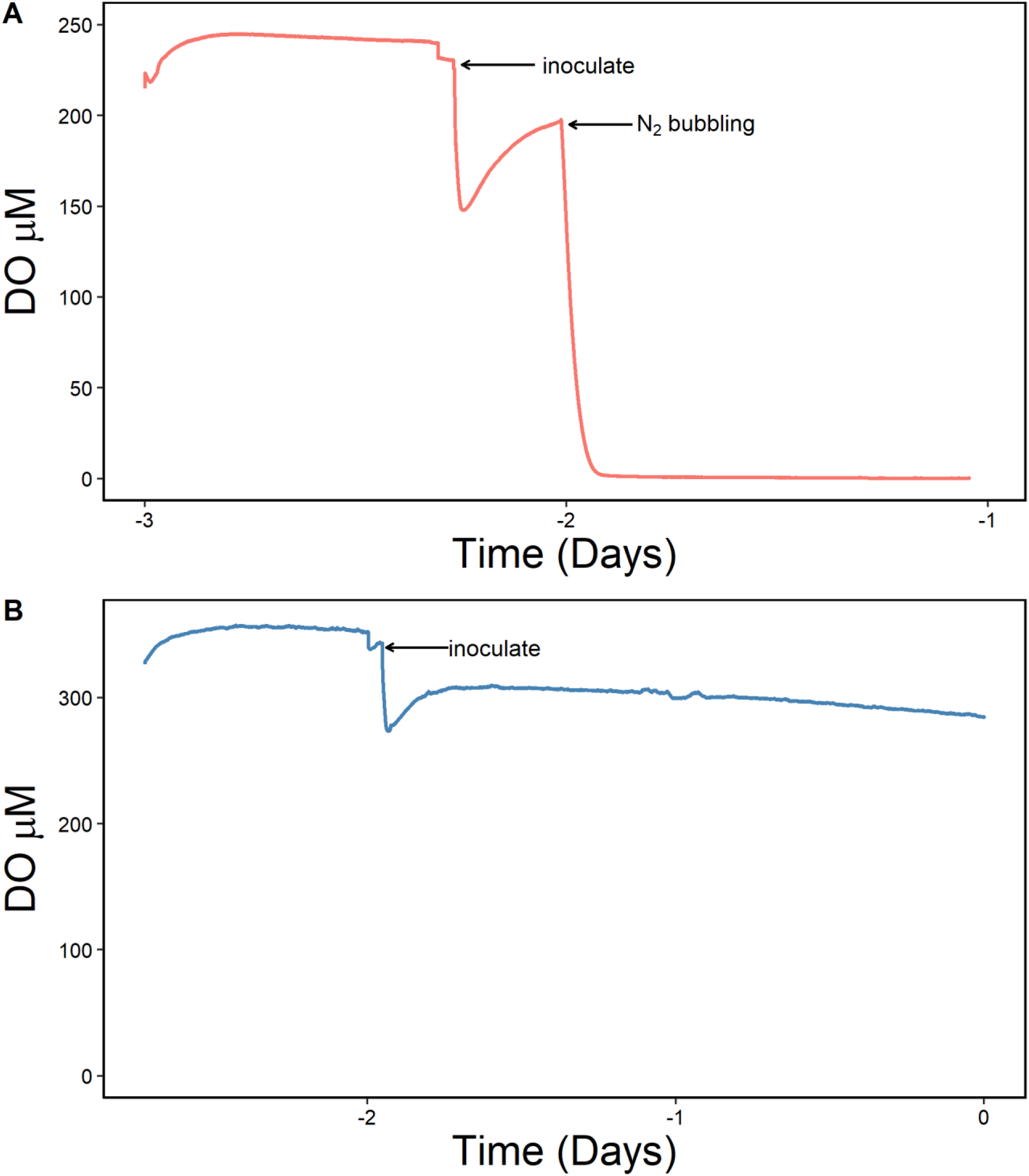
Dissolved Oxygen in Microaerobic and Passively Aerobic BES. Measurements of DO in BES under N_2_ bubbling (A) and passive aeration (B) with the small fiber optic probe. T=0 corresponds to addition of acetoin. DO concentration for N_2_ bubbled reactors ∼0.5-1.0 μM O_2_, and ∼300 μM O_2_ after inoculation.

## References

1. Rabaey K, Rozendal RA. 2010. Microbial electrosynthesis — revisiting the electrical route for microbial production. Nat Rev Microbiol 8:706–716.

2. Karthikeyan R, Singh R, Bose A. 2019. Microbial electron uptake in microbial electrosynthesis: a mini-review. J Ind Microbiol Biotechnol 46:1419–1426.

3. Choi O, Sang BI. 2016. Extracellular electron transfer from cathode to microbes: Application for biofuel production. Biotechnol Biofuels 9:1–14.

4. Joshi K, Kane AL, Kotloski NJ, Gralnick JA, Bond DR. 2019. Preventing Hydrogen Disposal Increases Electrode Utilization Efficiency by Shewanella oneidensis. Front Energy Res 7:95.

5. Xiong J, Chan D, Guo X, Chang F, Chen M, Wang Q, Song X, Wu C. 2020. Hydrogen production driven by formate oxidation in Shewanella oneidensis MR-1. Appl Microbiol Biotechnol 104:5579–5591.

6. Hirose A, Kouzuma A, Watanabe K. 2021. Hydrogen-dependent current generation and energy conservation by *Shewanella oneidensis* MR-1 in bioelectrochemical systems. J Biosci Bioeng 131:27–32.

7. Gong Z, Yu H, Zhang J, Li F, Song H. 2020. Microbial electro-fermentation for synthesis of chemicals and biofuels driven by bi-directional extracellular electron transfer. Synth Syst Biotechnol 5:304–313.

8. Prévoteau A, Carvajal-Arroyo JM, Ganigué R, Rabaey K. 2020. Microbial electrosynthesis from CO_2_: forever a promise? Curr Opin Biotechnol 62:48–57.

9. Bajracharya S, Ter Heijne A, Dominguez Benetton X, Vanbroekhoven K, Buisman CJN, Strik DPBTB, Pant D. 2015. Carbon dioxide reduction by mixed and pure cultures in microbial electrosynthesis using an assembly of graphite felt and stainless steel as a cathode. Bioresour Technol 195:14–24.

10. Harnisch F, Holtmann D, Batlle-Vilanova P, Berger C, Enzmann F, Geppert F, Gescher J, Halan B, Kerzenmacher S, Korth B, Krieg T, Liu D, Lütz S, Madjarov J, Puig S, Rabaey K, Rosa LFM, Rosenbaum MA, Rosenthal K, Schippers A, Schmid A, Schmitz LM, Schmitz S, Simonte F, Sleutels THJA, Sturm G, Sturm-Richter K, Tanne CK, ter Heijne A, Trschörtner J, Uhlig R, Vidakovic-Koch T. 2019. Bioelectrosynthesis.

11. Hengsbach JN, Sabel-Becker B, Ulber R, Holtmann D. 2022. Microbial electrosynthesis of methane and acetate—comparison of pure and mixed cultures. Appl Microbiol Biotechnol 106:4427–4443.

12. Gleizer S, Ben-Nissan R, Bar-On YM, Antonovsky N, Noor E, Zohar Y, Jona G, Krieger E, Shamshoum M, Bar-Even A, Milo R. 2019. Conversion of *Escherichia coli* to Generate All Biomass Carbon from CO_2_. Cell 179:1255–1263.

13. Herz E, Antonovsky N, Bar-On Y, Davidi D, Gleizer S, Prywes N, Noda-Garcia L, Lyn Frisch K, Zohar Y, Wernick DG, Savidor A, Barenholz U, Milo R. 2017. The genetic basis for the adaptation of *E. coli* to sugar synthesis from CO_2_. Nat Commun 8:1–10.

14. Antonovsky N, Gleizer S, Noor E, Zohar Y, Herz E, Barenholz U, Zelcbuch L, Amram S, Wides A, Tepper N, Davidi D, Bar-On Y, Bareia T, Wernick DG, Shani I, Malitsky S, Jona G, Bar-Even A, Milo R. 2016. Sugar Synthesis from CO_2_ in *Escherichia coli*. Cell 166:115–125.

15. Flamholz AI, Dugan E, Blikstad C, Gleizer S, Ben-nissan R, Amram S, Antonovsky N, Ravishankar S, Noor E. 2020. Functional reconstitution of a bacterial CO_2_ concentrating mechanism in *Escherichia coli*. Elife 2:1–30.

16. Coursolle D, Gralnick JA. 2010. Modularity of the Mtr respiratory pathway of *Shewanella oneidensis* strain MR-1. Mol Microbiol 77:995–1008.

17. Coursolle D, Gralnick JA. 2012. Reconstruction of extracellular respiratory pathways for iron(III) reduction in *Shewanella oneidensis* strain MR-1. Front Microbiol 3:56.

18. Nealson KH, Rowe AR. 2016. Electromicrobiology: realities, grand challenges, goals and predictions. Microb Biotechnol 9:595–600.

19. Ikeda S, Takamatsu Y, Tsuchiya M, Suga K, Tanaka Y, Kouzuma A, Watanabe K. 2021. *Shewanella oneidensis* MR-1 as a bacterial platform for electro-biotechnology. Essays Biochem 65:355–364.

20. Mitchell AC, Peterson L, Reardon CL, Reed SB, Culley DE, Romine MR, Geesey GG. 2012. Role of outer membrane c-type cytochromes MtrC and OmcA in *Shewanella oneidensis* MR-1 cell production, accumulation, and detachment during respiration on hematite. Geobiology 10:355– 370.

21. Fredrickson JK, Romine MF, Beliaev AS, Auchtung JM, Driscoll ME, Gardner TS, Nealson KH, Osterman AL, Pinchuk G, Reed JL, Rodionov DA, Rodrigues JLM, Saffarini DA, Serres MH, Spormann AM, Zhulin IB, Tiedje JM. 2008. Towards environmental systems biology of *Shewanella*. Nat Rev Microbiol 6:592–603.

22. Marritt SJ, Lowe TG, Bye J, McMillan DGG, Shi L, Fredrickson J, Zachara J, Richardson DJ, Cheesman MR, Jeuken LJC, Butt JN. 2012. A functional description of CymA, an electron-transfer hub supporting anaerobic respiratory flexibility in *Shewanella*. Biochemical Journal 444:465–474.

23. Sturm G, Richter K, Doetsch A, Heide H, Louro RO, Gescher J. 2015. A dynamic periplasmic electron transfer network enables respiratory flexibility beyond a thermodynamic regulatory regime. ISME Journal 9:1802–1811.

24. McMillan DGG, Marritt SJ, Butt JN, Jeuken LJC. 2012. Menaquinone-7 is specific cofactor in tetraheme quinol dehydrogenase CymA. Journal of Biological Chemistry 287:14215–14225.

25. Ross DE, Flynn JM, Baron DB, Gralnick JA, Bond DR. 2011. Towards electrosynthesis in *Shewanella*: Energetics of reversing the Mtr pathway for reductive metabolism. PLoS One 6:1–9.

26. Rowe AR, Salimijazi F, Trutschel L, Sackett J, Adesina O, Anzai I, Kugelmass LH, Baym MH, Barstow B. 2021. Identification of a pathway for electron uptake in *Shewanella oneidensis*. Commun Biol 4:1–10.

27. Edwards MJ, White GF, Lockwood CW, Lawes MC, Martel A, Harris G, Scott DJ, Richardson DJ, Butt JN, Clarke TA. 2018. Structural modeling of an outer membrane electron conduit from a metal-reducing bacterium suggests electron transfer via periplasmic redox partners. Journal of Biological Chemistry 293:8103–8112.

28. Tefft NM, TerAvest MA. 2019. Reversing an Extracellular Electron Transfer Pathway for Electrode-Driven Acetoin Reduction. ACS Synth Biol 8:1590–1600.

29. Tefft NM, Ford K, TerAvest MA. 2022. NADH dehydrogenases drive inward electron transfer in *Shewanella oneidensis* MR-1. Microb Biotechnol 00:1–9.

30. Rabaey K, Read ST, Clauwaert P, Freguia S, Bond PL, Blackall LL, Keller J. 2008. Cathodic oxygen reduction catalyzed by bacteria in microbial fuel cells. ISME Journal 2:519–527.

31. Rosenbaum M, Aulenta F, Villano M, Angenent LT. 2011. Cathodes as electron donors for microbial metabolism: Which extracellular electron transfer mechanisms are involved? Bioresour Technol 102:324–333.

32. Rowe AR, Rajeev P, Jain A, Pirbadian S, Okamoto A, Gralnick JA, El-Naggar MY, Nealson KH. 2018. Tracking electron uptake from a cathode into *Shewanella* cells: Implications for energy acquisition from solid-substrate electron donors. mBio 9:1–19.

33. Tefft NM, Ford K, TerAvest MA. 2022. NADH dehydrogenases drive inward electron transfer in Shewanella oneidensis MR-1. Microb Biotechnol 00:1–9.

34. Duhl KL, Tefft NM, TerAvest MA. 2018. *Shewanella oneidensis* MR-1 utilizes both sodium-and proton-pumping NADH dehydrogenases during aerobic growth. Appl Environ Microbiol 84:1–12.

35. Duhl KL, TerAvest MA. 2019. *Shewanella oneidensis* NADH Dehydrogenase Mutants Exhibit an Amino Acid Synthesis Defect. Front Energy Res 7:1–12.

36. Madsen CS, TerAvest MA. 2019. NADH dehydrogenases Nuo and Nqr1 contribute to extracellular electron transfer by *Shewanella oneidensis* MR-1 in bioelectrochemical systems. Sci Rep 9:1–6.

37. Chen H, Luo Q, Yin J, Gao T, Gao H. 2015. Evidence for the requirement of CydX in function but not assembly of the cytochrome *bd* oxidase in *Shewanella oneidensis*. Biochim Biophys Acta Gen Subj 1850:318–328.

38. Yin J, Meng Q, Fu H, Gao H. 2016. Reduced expression of cytochrome oxidases largely explains cAMP inhibition of aerobic growth in *Shewanella oneidensis*. Sci Rep 6:1–11.

39. Le Laz S, Kpebe A, Bauzan M, Lignon S, Rousset M, Brugna M. 2014. A biochemical approach to study the role of the terminal oxidases in aerobic respiration Shewanella oneidensis MR-1. PLoS One 9:1–10.

40. Mortimer CH. 1956. The oxygen content of air-saturated fresh waters, and aids in calculating percentage saturation. SIL Communications, 1953-1996 6:1–20.

41. Zhou G, Yin J, Chen H, Hua Y, Sun L, Gao H. 2013. Combined effect of loss of the *caa_3_* oxidase and Crp regulation drives *Shewanella* to thrive in redox-stratified environments. ISME Journal 7:1752–1763.

42. Norman MP, Edwards MJ, White GF, Burton JAJ, Louro RO, Paquete CM, Clarke A. 2023. A Cysteine Pair Controls Flavin Reduction by Extracellular Cytochromes during Anoxic/Oxic Environmental Transitions. mBio 14:1–12.

43. Burek BO, Bormann S, Hollmann F, Bloh JZ, Holtmann D. 2019. Hydrogen peroxide driven biocatalysis. Green Chemistry 21:3232–3249.

44. Massey Vincent. 1994. Activation of Molecular Oxygen by Flavins and Flavoproteins. Journal of Biological Chemistry 269:22459–22462.

45. Roncel M, Navarro JA, Rosa FFD la, Rosa MAD la. 1984. Flavin-Mediated Production of Hydrogen Peroxide in Photoelectrochemical Cells. Photochem Photobiol 40:395–398.

46. Zhou ZY, Zhang HT, Shi Y. 2004. Theoretical study of the interactions between 1,3-butanediol and hydrogen peroxide. Journal of Physical Chemistry A 108:6520–6526.

47. Sato K, Aoki M, Takagi J, Noyori R. 1998. Organic solvent-and halide-free oxidation of alcohols with aqueous hydrogen peroxide. Chemtracts 11:629–631.

48. Hartshorne RS, Reardon CL, Ross D, Nuester J, Clarke TA, Gates AJ, Mills PC, Fredrickson JK, Zachara JM, Shi L, Beliaev AS, Marshall MJ, Tien M, Brantley S, Butt JN, Richardson DJ. 2009. Characterization of an electron conduit between bacteria and the extracellular environment. Proceedings of the National Academy of Sciences 106:22169–22174.

49. Harada E, Kumagai J, Ozawa K, Imabayashi S, Tsapin AS, Nealson KH, Meyer TE, Cusanovich MA, Akutsu H. 2002. A directional electron transfer regulator based on heme-chain architecture in the small tetraheme cytochrome *c* from *Shewanella oneidensis*. FEBS Lett 532:333–337.

50. Pessanha M, Rothery EL, Miles CS, Reid GA, Chapman SK, Louro RO, Turner DL, Salgueiro CA, Xavier A V. 2009. Tuning of functional heme reduction potentials in *Shewanella* fumarate reductases. Biochim Biophys Acta Bioenerg 1787:113–120.

51. White GF, Shi Z, Shi L, Wang Z, Dohnalkova AC, Marshall MJ, Fredrickson JK, Zachara JM, Butt JN, Richardson DJ, Clarke TA. 2013. Rapid electron exchange between surface-exposed bacterial cytochromes and Fe(III) minerals. Proc Natl Acad Sci U S A 110:6346–6351.

52. Firer-Sherwood M, Pulcu GS, Elliott SJ. 2008. Electrochemical interrogations of the Mtr cytochromes from *Shewanella*: Opening a potential window. Journal of Biological Inorganic Chemistry 13:849–854.

53. Frantz NL, Brakoniecki G, Chen D, Proshlyakov DA. 2021. Assessment of the maximal activity of complex IV in the inner mitochondrial membrane by tandem electrochemistry and respirometry. Anal Chem 93:1360–1368.

54. Wickham H. 2016. ggplot2: Elegant Graphics for Data Analysis. Springer-Verlag New York. ISBN 978–3-319-24277-4, https://ggplot2.tidyverse.org.

55. Wickham H, François R, Henry L. 2020. dplyr: A Grammar of Data Manipulation. https://dplyr.tidyverse.org, https://github.com/tidyverse/dplyr.

56. Kassambara A. 2023. ggpubr:“ggplot2” based publication ready plots. https://CRANR-project.org/package=ggpubr 438.

57. Wickham H, Wickham MH. 2011. The Split-Apply-Combine Strategy for Data Analysis. J Stat Softw 40:1–29.

58. Dowle M, Srinivasan A, Gorecki J, Chirico M, Stetsenko P, Short T, Lianoglou S, Antonyan E, Bonsch M, Parsonage H. 2019. data.table: Extension of “data.frame.” https://CRANR-project.org/package=data.table596.

59. Wickham H. 2022. stringr: Simple, Consistent Wrappers for Common String Operations. http://stringrtidyverse.org, https://githubcom/tidyverse/stringr.org.

60. Sprouffske K, Wagner A. 2016. Growthcurver: an R package for obtaining interpretable metrics from microbial growth curves. BMC Bioinformatics 17:1–4.

